# Frequent and asymmetric cell division in endosymbiotic bacteria of cockroaches

**DOI:** 10.1101/2024.05.17.594780

**Authors:** Tomohito Noda, Masaki Mizutani, Toshiyuki Harumoto, Tatsuya Katsuno, Ryuichi Koga, Takema Fukatsu

**Author notes:** Correspondence: Takema Fukatsu. These authors have contributed equally to this work (Tomohito Noda, Masaki Mizutani).

## Abstract

Many insects are obligatorily associated with and dependent on specific microbial species as essential mutualistic partners. In the host insects, such microbial mutualists are usually maintained in specialized cells or organs, called bacteriocytes or symbiotic organs. Hence, potentially exponential microbial growth cannot be realized but must be strongly constrained by spatial and resource limitations within the host cells or tissues. How such endosymbiotic bacteria grow, divide and proliferate is important for understanding the interactions and dynamics underpinning intimate host-microbe symbiotic associations. Here we report that *Blattabacterium*, the ancient and essential endosymbiont of cockroaches, exhibits unexpectedly high rates of cell division (20-58%) and, in addition, the cell division is asymmetric (average asymmetry index > 1.5) when isolated from the German cockroach *Blattella germanica*. The asymmetric division of endosymbiont cells at high frequencies was observed irrespective of host tissues (fat bodies vs. ovaries) or developmental stages (adults vs. nymphs vs. embryos) of *B. germanica*, and also observed in several different cockroach species. By contrast, such asymmetric and frequent cell division was observed neither in *Buchnera*, the obligatory bacterial endosymbiont of aphids, nor in *Pantoea*, the obligatory bacterial gut symbiont of stinkbugs. Comparative genomics of cell division-related genes uncovered that the *Blattabacterium* genome lacks the Min system genes that determine the cell division plane, which may be relevant to the asymmetric cell division. These observations combined with comparative symbiont genomics provide insight into what processes and regulations may underpin the growth, division and proliferation of such bacterial mutualists continuously constrained under within-host conditions.

**IMPORTANCE:** Diverse insects are dependent on specific bacterial mutualists for their survival and reproduction. Due to the long-lasting coevolutionary history, such symbiotic bacteria tend to exhibit degenerative genomes and suffer uncultivability. Because of their microbiological fastidiousness, the cell division patterns of such uncultivable symbiotic bacteria have been poorly described. Here, using fine microscopic and quantitative morphometric approaches, we report that, although bacterial cell division usually proceeds through symmetric binary fission, *Blattabacterium*, the ancient and essential endosymbiont of cockroaches, exhibits frequent and asymmetric cell division. Such peculiar cell division patterns were not observed with other uncultivable essential symbiotic bacteria of aphids and stinkbugs. Gene repertoire analysis revealed that the molecular machineries for regulating the bacterial cell division plane are lost in the *Blattabacterium* genome, suggesting the possibility that the general trend toward the reductive genome evolution of symbiotic bacteria may underpin their bizarre cytological/morphological traits.

## INTRODUCTION

Cell division is a fundamental process for all cellular organisms. In many bacteria, the cell division proceeds through a process called binary fission. Typically, each rod-shaped bacterial cell experiences growth, septation, and division during the cell cycle, under which such cytological processes as DNA replication, Z-ring formation, septum formation and cell division proceed. Molecular mechanisms and machineries underpinning the bacterial cell division have been extensively investigated in several model bacteria like *Escherichia coli*, *Bacillus subtilis* and *Caulobacter crescentus* (1–3). On the other hand, reflecting the enormous diversity of bacteria, striking diversity has been described not only in bacterial cellular shapes (coccal, elongate, spiral, crescent, filamentous, pleomorphic, etc.) but also in modes of bacterial cell division (asymmetric fission, endospore formation, budding, baeocyte formation, arthrospore formation, etc.), whose underlying mechanisms are still to be investigated (4–6).

Many insects are obligatorily associated with and dependent on specific symbiotic bacteria as essential mutualistic partners. Such microbial symbionts are essential for growth, survival, and reproduction of their hosts by supplying essential nutrients (7,8), assisting food digestion (9,10), or defense against natural enemies (11,12). In these symbiotic associations, the host insects often develop specialized cells, tissues and organs for harboring the symbiotic bacteria. Some insects maintain their symbiotic bacteria extracellularly within the inner cavity of specialized structures associated with their alimentary tract, called crypts or gastric caeca, as in stinkbugs and leaf beetles (13–15). Other insects retain their symbiotic bacteria intracellularly within the cytoplasm of specialized cells and organs, called bacteriocytes and bacteriomes, as in aphids and cockroaches (13,16,17). During the intimate host-symbiont association over evolutionary time, the symbiotic bacteria tend to experience degenerative genome evolution wherein many bacterial genes needed for a free-living lifestyle yet dispensable for within-host lifestyle are eroded and lost, thereby resulting in a variety of peculiar microbial traits such as reduced genome size, uncultivability, strange cell morphology, etc. (7,18,19). When taken out of the host insects, such symbiotic bacteria can neither survive nor grow, often showing a variety of cell shapes deviated from typical rod, coccal or tubular ones to very large, very long, rosette-like, pleomorphic or deformed ones (13,20–24), which may entail amplified genome copies per cell called polyploidy (25–27).

The strict within-host lifestyle of the obligatory symbiotic bacteria is expected to constrain the potentially exponential microbial growth due to spatial and resource limitations, which may affect their cell division, proliferation and cell cycle patterns. However, few previous studies have investigated such cell biological aspects of the obligatory symbiotic bacteria of insects. Although there are some microscopic observations that identified dividing cells of such uncultivable symbiotic bacteria of insects (see 13), these observations are neither systematic nor quantitative but are mostly mere snapshot images. Although there are a number of reports on the population dynamics of such uncultivable symbiotic bacteria (ex. 28-32), they are usually based on quantitative PCR detection with little cytological information.

In this study, we report that *Blattabacterium*, the ancient and essential endosymbiont of cockroaches involved in nitrogen recycling and amino acid provisioning (33–36), exhibits unexpectedly high rates of cell division (20-58%) and, in addition, the cell division is asymmetric (average asymmetry index > 1.5) when isolated from the host German cockroach *Blattella germanica*. We conducted quantitative comparisons of cell size, cell division rate and asymmetry level of the symbiotic bacteria between different tissues and developmental stages of *B. germanica*, and also between different cockroach species. Furthermore, we extended the comparative analyses to the obligatory symbiotic bacteria of other insects, *Buchnera* of an aphid and *Pantoea* of a stinkbug, and moreover to the cultivable model bacterium *Escherichia coli* using morphometric and comparative genomic approaches.

## RESULTS

### Frequent and asymmetric cell division in *Blattabacterium* endosymbiont of *B. germanica*

From five adult insects of *B. germanica* (Fig. 1A), fat bodies were dissected out (Fig. 1B), homogenized and diluted, and observed under a phase-contrast microscope, by which *Blattabacterium* endosymbionts were visualized as rod-shaped bacterial cells (Fig. 1C-I). Notably, we found that the bacterial cells exhibited asymmetric division at strikingly high rates (20.5%–41.5% in 200 bacterial cells per insect) (Fig. 1J-O; Table S1). Electron microscopic observations of bacteriocytes in the adult fat bodies confirmed the presence of dividing bacterial cells in the process of asymmetric septum formation (Fig. 2A, B). The dividing bacterial cells (7.67 ± 1.76 μm, N = 50) were longer than the non-dividing bacterial cells (5.54 ± 1.35 μm, N = 50), where the difference was highly significant statistically (*P* = 1.0 x 10^-9^) (Fig. 1P; Table S1), indicating that the dividing bacterial cells are well-grown cells in division. The asymmetry indices of the dividing bacterial cells, which were calculated by the length of the longer daughter cell divided by the length of the shorter daughter cell, were conspicuously deviated from 1, with mean and median being 1.79 and 1.59, respectively (Fig. 1Q; Table S2). These observations uncovered that *Blattabacterium* endosymbiont cells harbored in the bacteriocytes within the fat bodies of *B. germanica* exhibit asymmetric division at high frequencies.

**FIG 1.**
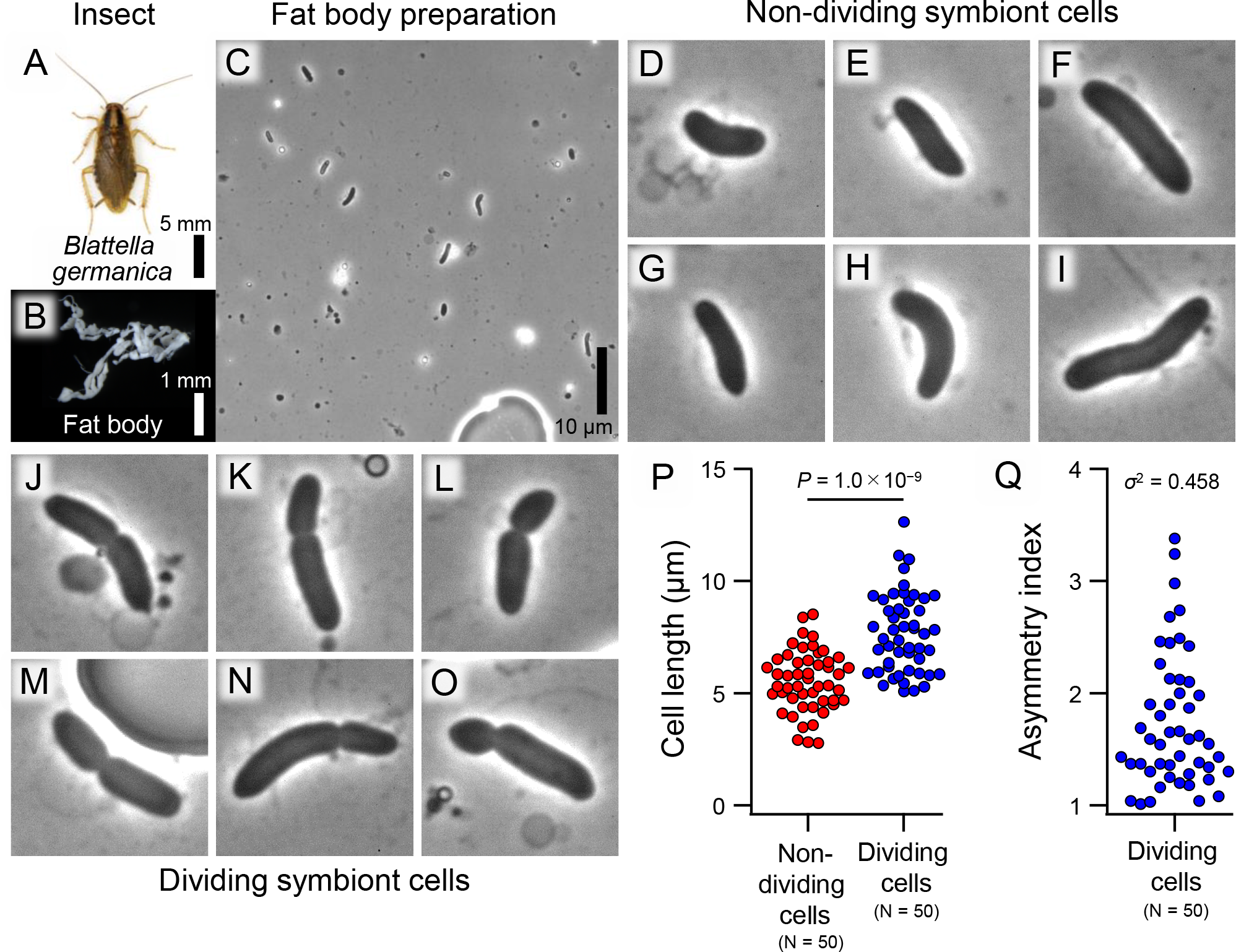
Morphology of *Blattabacterium* symbiont cells prepared from fat bodies of the adult German cockroach *B. germanica.* (**A**) An adult insect. (**B**) Dissected fat bodies. (**C-O**) Phase-contrast microscopic images of isolated symbiont cells. (**C**) A low magnification image. (**D-I**) Magnified images of non-dividing symbiont cells. (**J-O**) Magnified images of dividing symbiont cells. (**P**) Comparison of lengths of non-dividing and dividing symbiont cells. The difference is significant statistically (Student’s *t*-test: *P* = 1.0 x 10^-9^). Also see Table S1. (**Q**) Asymmetry indices of dividing symbiont cells, which are defined as length of the longer daughter cell divided by length of the shorter daughter cell. Also see Table S2. Panels (**D-O**) are 10 μm x 10 μm squares.

**FIG 2.**
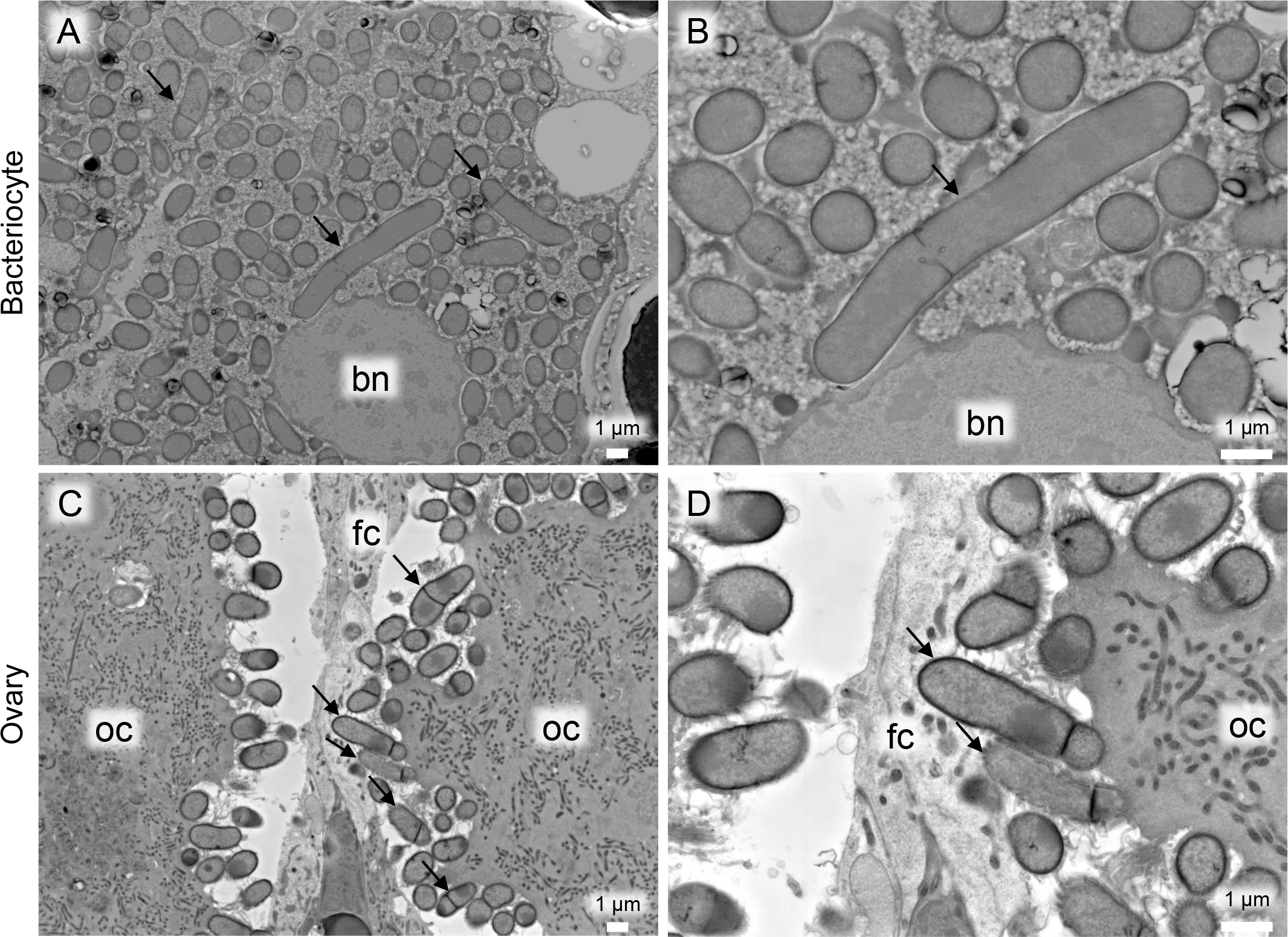
Backscattered electron scanning electron microscopic images of *Blattabacterium* symbiont cells of *B. germanica*. (**A, B**) Symbiont cells packed in a bacteriocyte, which is embedded within the fat body. (**C, D**) Symbiont cells associated with developing oocytes in the ovary. Arrows indicate symbiont cells in asymmetric division. Abbreviations: bn, bacteriocyte nucleus; oc, oocyte; fc, follicle cell.

### Frequent and asymmetric cell division in *Blattabacterium* endosymbiont from different tissues and developmental stages of *B. germanica*

When dissected ovaries of *B. germanica* were homogenized, diluted and inspected under a phase-contrast microscope, *Blattabacterium* endosymbionts were observed as rod-shaped bacterial cells with frequent asymmetric division at a rate of 30.0% (Fig. 3A-J; Table S1). Electron microscopic observations of the adult ovaries confirmed the presence of bacterial cells in asymmetric division (Fig. 2C, D). Similarly, *Blattabacterium* endosymbiont cells exhibited asymmetric division at high rates, 58.0% for ovaries dissected from 2^nd^ instar nymphs and 47.0% for embryos dissected from oothecae (Fig. 3K-e; Table S1). Size comparisons of the dividing and non-dividing bacterial cells revealed that (i) the dividing cells were consistently and significantly larger in size than the non-dividing cells for each of adult ovaries, embryos and nymphal ovaries, (ii) the non-dividing cells were not different in size among adult ovaries, nymphal ovaries and embryos, and (iii) the dividing cells were also not different in size among adult ovaries, nymphal ovaries and embryos (Fig. 3f; Table S2). The asymmetry indices of the dividing bacterial cells were consistently deviated from 1, with mean and median being 1.56 and 1.42 for adult ovaries, 1.68 and 1.45 for nymphal ovaries, and 1.67 and 1.46 for embryos (Fig. 3g; Table S2). These results indicated that *Blattabacterium* endosymbiont cells consistently exhibit asymmetric division at high frequencies irrespective of host tissues (fat bodies vs. ovaries) or developmental stages (adults vs. nymphs vs. embryos).

**FIG 3.**
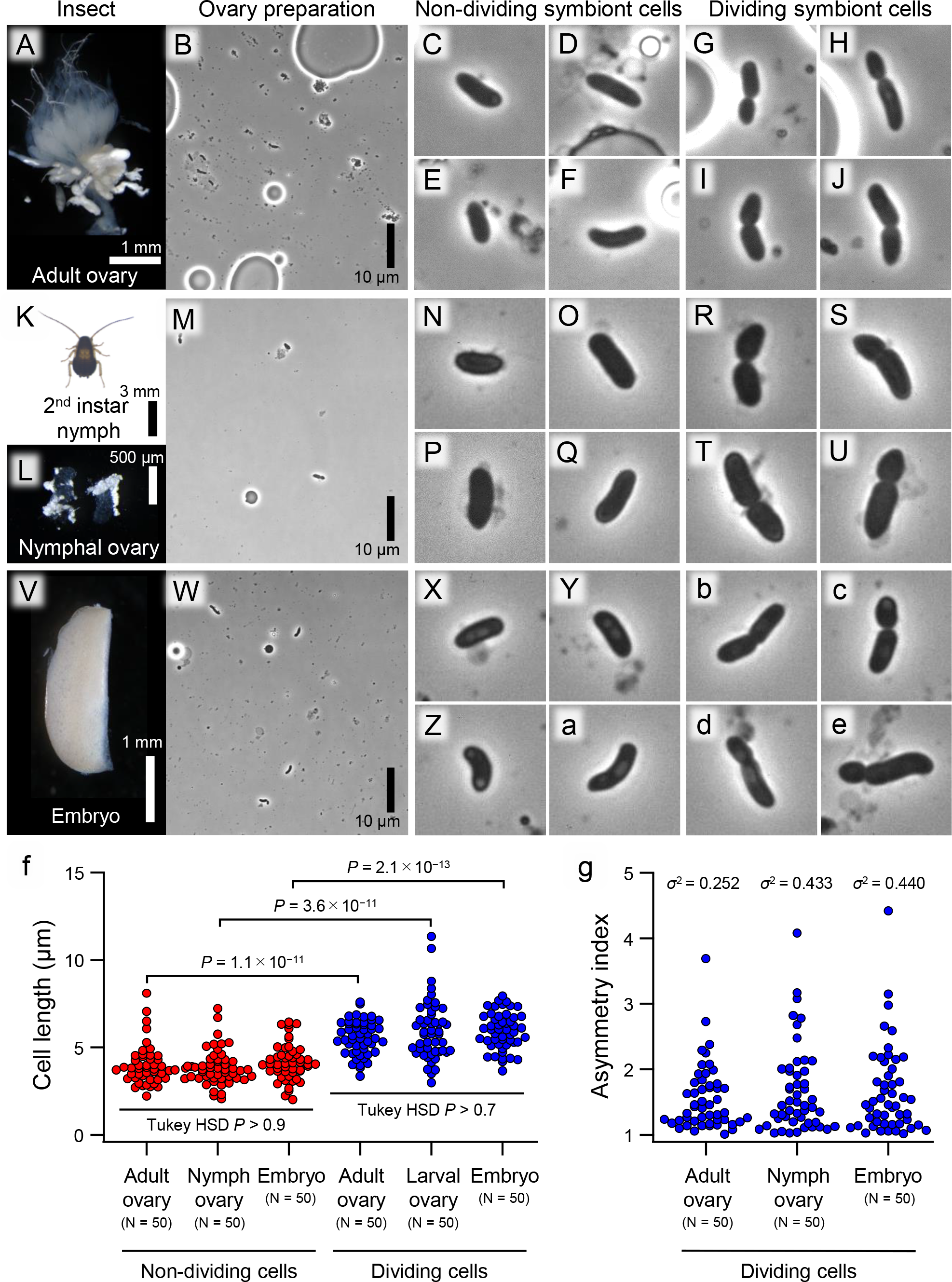
Morphology of *Blattabacterium* symbiont cells prepared from adult ovaries, nymphal ovaries and embryos of the German cockroach *B. germanica.* (**A-J**) Data from ovaries dissected from adult insects. (**A**) Dissected ovaries. (**B-J**) Phase-contrast microscopic images of isolated symbiont cells. (**B**) A low magnification image. (**C-F**) Magnified images of non-dividing symbiont cells. (**G-J**) Magnified images of dividing symbiont cells. (**K-U**) Data from ovaries dissected from 2^nd^ instar nymphs. (**K**) A 2^nd^ instar nymph. (**L**) Dissected ovaries. (**M-U**) Phase-contrast microscopic images of isolated symbiont cells. (**M**) A low magnification image. (**N-Q**) Magnified images of non-dividing symbiont cells. (**R-U**) Magnified images of dividing symbiont cells. (**V-e**) Data from embryos dissected from oothecae. (**V**) An embryo. (**W-e**) Phase-contrast microscopic images of isolated symbiont cells. (**W**) A low magnification image. (**X-a**) Magnified images of non-dividing symbiont cells. (**b-e**) Magnified images of dividing symbiont cells. (**f**) Comparisons of lengths of non-dividing symbiont cells from adult ovaries, nymphal ovaries and embryos, and also lengths of dividing symbiont cells from adult ovaries, nymphal ovaries and embryos. The differences within the categories, namely non-dividing cells and dividing cells respectively, are not significant statistically (Tukey HSD test: *P* > 0.9, *P* > 0.7), whereas the differences between the categories for each material, namely adult ovaries, nymphal ovaries or embryos, are all statistically significant (Student’s *t*-test: *P* = 1.1 x 10^−11^, *P* = 3.6 x 10^−11^, *P* = 2.1 x 10^−13^). Also see Table S1. (**g**) Asymmetry indices of dividing symbiont cells, which are defined as length of the longer daughter cell divided by length of the shorter daughter cell, for adult ovaries, nymphal ovaries and embryos. Also see Table S2. Panels (**C-J**), (**N-U**) and (**X-e**) are 10 μm x 10 μm squares.

### Frequent and asymmetric cell division in *Blattabacterium* endosymbionts of different cockroach species

Does the asymmetric division of *Blattabacterium* endosymbiont cells at high frequencies also occur in other cockroach species? To address this question, we conducted similar microscopic observations on fat body preparations of *Periplaneta japonica* (Fig. 4A-J), *Eucorydia yasumatsui* (Fig. 4K-T) and *Pycnoscelus indicus* (Fig. 4U-d). In all the cockroach species, asymmetric division of endosymbiont cells were observed (Fig. 4G-J, Q-T and a-d), but the frequencies were different between the species: 34.5% for *P. japonica* and 24.0% for *E. yasumatsui* were high, but 8.0% for *P. indicus* was not (Table S1). Size comparisons of the dividing and non-dividing bacterial cells showed that the dividing cells were significantly larger in size than the non-dividing cells in *P. japonica* and *E. yasumatsui*, but not in *P. indicus* (Fig. 4e; Table S2). The asymmetry indices of the dividing bacterial cells were consistently deviated from 1, with mean and median being 1.49 and 1.39 for *P. japonica*, 1.72 and 1.46 for *E. yasumatsui*, and 1.46 and 1.28 for *P. indicus* (Fig. 4f; Table S2). These results indicated that the asymmetric division of *Blattabacterium* endosymbiont cells is widely observed among different cockroach species, but frequency of dividing bacterial cells may vary among the species.

**FIG 4.**
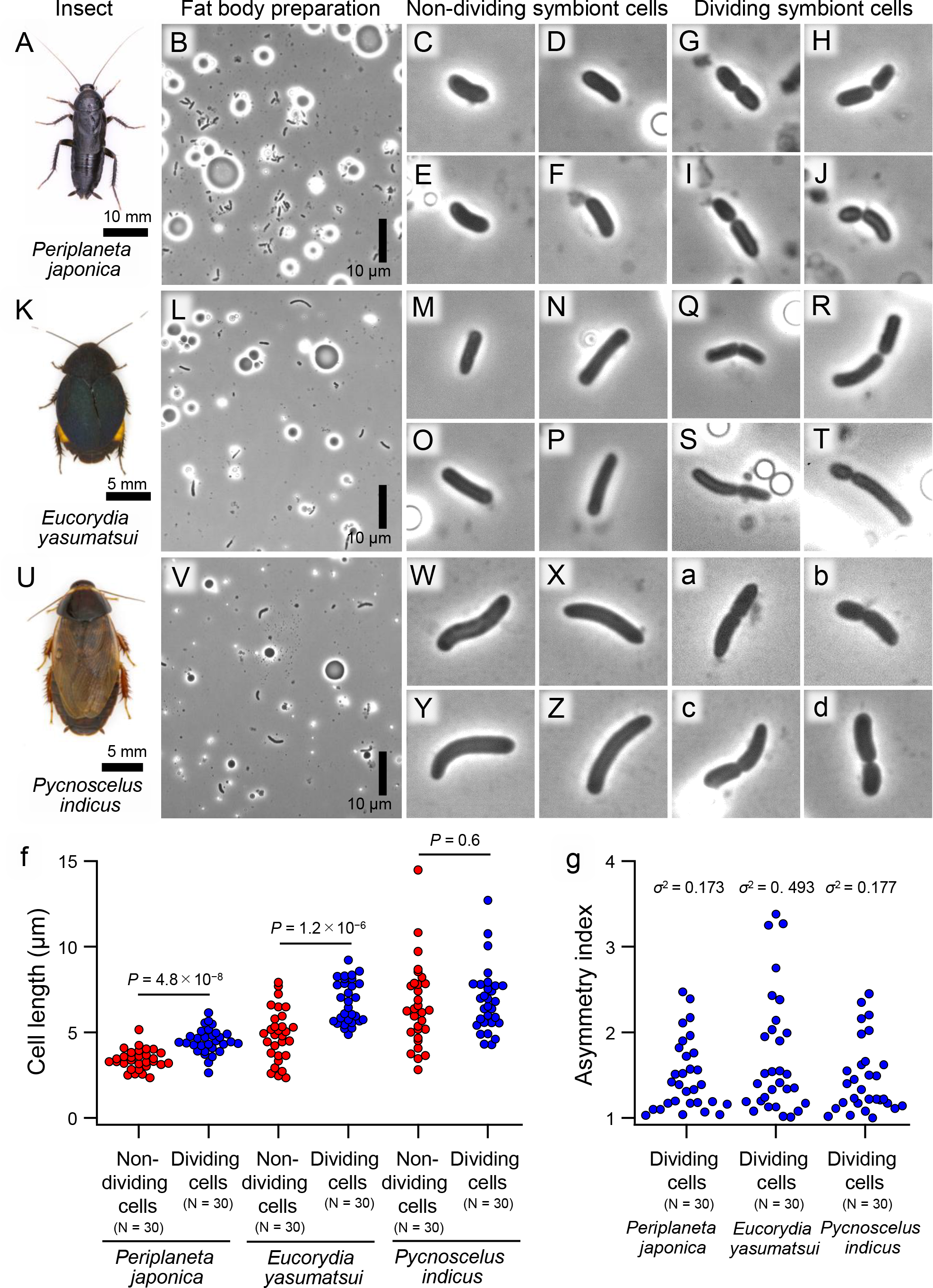
Morphology of *Blattabacterium* symbiont cells prepared from different cockroach species. (**A-J**) Data from the Japanese cockroach *Periplaneta japonica*. (**A**) An adult insect. (**B-J**) Phase-contrast microscopic images of isolated symbiont cells. (**B**) A low magnification image. (**C-F**) Magnified images of non-dividing symbiont cells. (**G-J**). Magnified images of dividing symbiont cells. (**K-T**) Data from the azure cockroach *Eucorydia yasumatsui*. (**K**) An adult insect. (**L-T**) Phase-contrast microscopic images of isolated symbiont cells. (**L**) A low magnification image. (**M-P**) Magnified images of non-dividing symbiont cells. (**Q-T**). Magnified images of dividing symbiont cells. (**U-d**) Data from the Indian cockroach *Pycnoscelus indicus*. (**U**) An adult insect. (**V-d**) Phase-contrast microscopic images of isolated symbiont cells. (**V**) A low magnification image. (**W-Z**) Magnified images of non-dividing symbiont cells. (**a-d**). Magnified images of dividing symbiont cells. (**e**) Comparisons of lengths of non-dividing and dividing symbiont cells from *P. japonica*, *E. yasumatsui* and *P. indicus*. The differences between non-dividing and dividing symbiont cells are significant statistically for *P. japonica* and *E. yasumatsui* (Student’s *t*-test: *P* = 4.8 x 10^−8^, *P* = 1.2 x 10^−6^), whereas no significant difference was detected for *P. inducus* (Student’s *t*-test: *P* = 0.6). Also see Table S1. (**f**) Asymmetry indices of dividing symbiont cells, which are defined as length of the longer daughter cell divided by length of the shorter daughter cell, for *P. japonica*, *E. yasumatsui* and *P. indica.* Also see Table S2. Panels (**C-J**), (**M-T**) and (**W-d**) are 10 μm x 10 μm squares.

### No asymmetric cell division in obligatory endosymbiont of aphid and obligatory gut symbiont of stinkbug

Is such asymmetric division of endosymbiont cells also observed in other insect-microbe symbiotic systems? To address this question, we conducted similar microscopic observations on bacteriocyte preparations of the aphid *Acyrthosiphonn pisum* associated with an obligatory bacterial endosymbiont *Buchnera* (Fig. 5A-K) and midgut symbiotic organ preparations of the stinkbug *Plautia stali* associated with an obligatory bacterial gut symbiont *Pantoea* (Fig. 5L-V). In both the species, the bacterial cells exhibited binary and equal cell division (Fig. 5H-K, S-V), whereas frequencies of dividing cells were moderately high (12.5%) for the aphid endosymbiont *Buchnera* and very low (5.0%) for the stinkbug gut symbiont *Pantoea* (Table S1). Size comparisons of dividing and non-dividing bacterial cells showed that the dividing cells were significantly larger in size than the non-dividing cells in both the aphid endosymbiont *Buchnera* and the stinkbug gut symbiont *Pantoea* (Fig. 5W; Table S2), confirming that the dividing bacterial cells are well-grown cells in division. The asymmetry indices of the dividing bacterial cells were almost 1, reflecting the equal cell division of the aphid endosymbiont *Buchnera* and the stinkbug gut symbiont *Pantoea* (Fig. 5X; Table S2).

**FIG 5.**
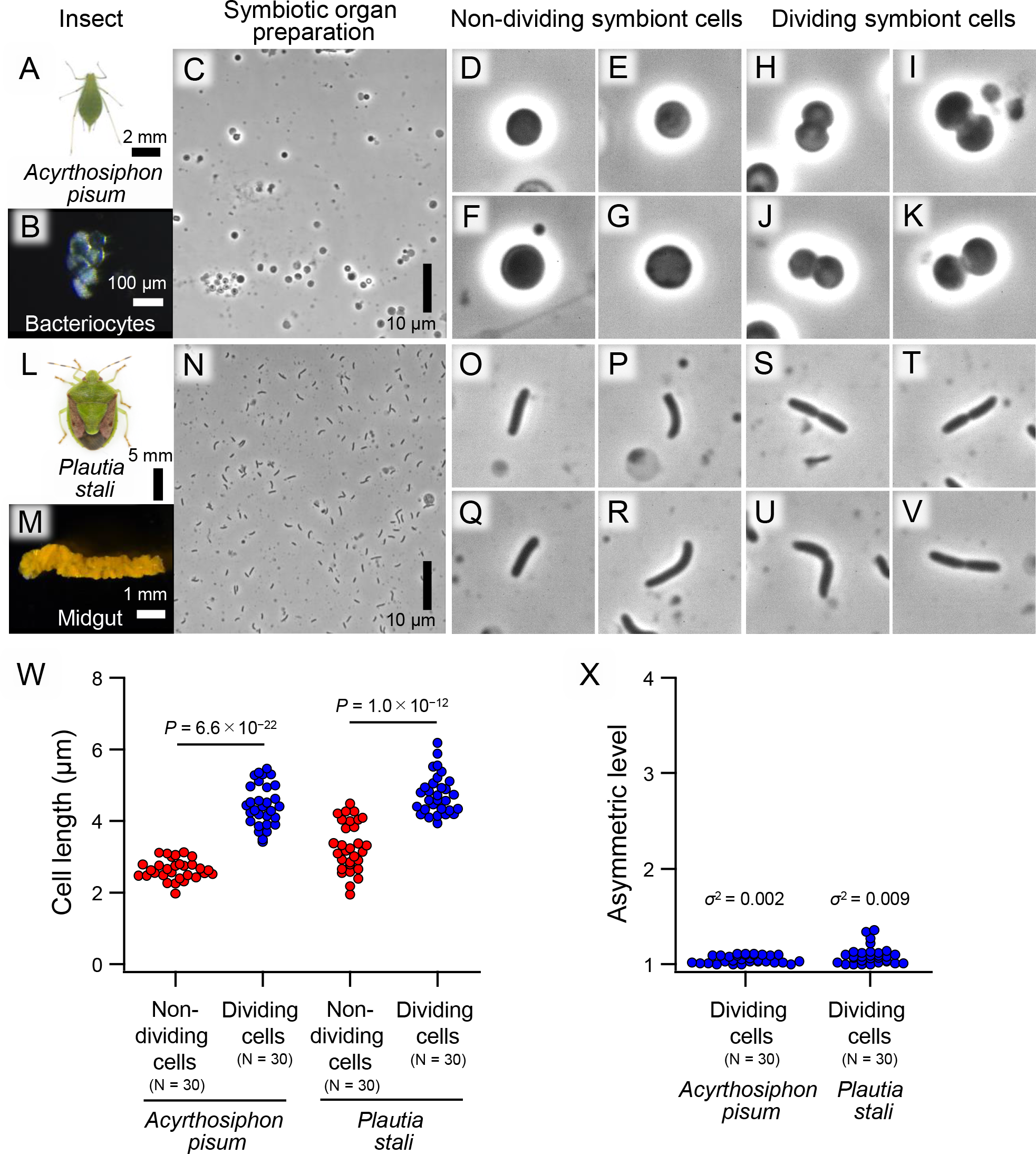
Morphology of symbiont cells from the pea aphid *Acyrthosiphon pisum* and the brown-winged stinkbug *Plautia stali*. (**A-K**) Data of *Buchnera* endosymbiont cells prepared from bacteriocytes of *A. pisum*. (**A**) An adult insect. (**B**) Dissected bacteriocytes. (**C-K**) Phase-contrast microscopic images of isolated symbiont cells. (**C**) A low magnification image. (**D-G**) Magnified images of non-dividing symbiont cells. (**H-K**). Magnified images of dividing symbiont cells. (**L-V**) Data of *Pantoea* gut symbiont cells prepared from the midgut symbiotic organ of *P. stali*. (**A**) An adult insect. (**M**) A dissected midgut symbiotic organ. (**N-V**) Phase-contrast microscopic images of isolated symbiont cells. (**N**) A low magnification image. (**O-R**) Magnified images of non-dividing symbiont cells. (**S-V**). Magnified images of dividing symbiont cells. (**W**) Comparisons of lengths of non-dividing and dividing symbiont cells. The differences are significant statistically for both *A. pisum* and *P. stali* (Student’s *t*-test: *P* = 6.6 x 10^-22^*, P* = 1.0 x 10^-12^). (**X**) Asymmetry indices of dividing symbiont cells, which are defined as length of the longer daughter cell divided by length of the shorter daughter cell, for *A. pisum* and *P. stali*. Also see Table S2. Panels (**D-K**) and (**O-V**) are 10 μm x 10 μm squares.

### Statistics and notes

Table S3 shows two-way ANOVA results on cell length comparisons between dividing symbiont cells and non-dividing symbiont cells from different tissues of *B. germanica*, which showed that both cell states (dividing vs. non-dividing) and tissue types (adult fat body vs. adult ovary vs. nymphal ovary vs. embryo) have statistically significant effects on the cell length of *Blattabacterium* (both *P* < 0.001). Table S4 summarizes *P* values of the post-hoc Tukey HSD test comparing the cell lengths of dividing symbiont cells and non-dividing symbiont cells from different tissues of *B. germanica*. We note that *Blattabacterium* cell length was significantly longer in those from adult fat bodies than in those from adult ovaries, nymphal ovaries and embryos of *B. germanica* irrespective of dividing or non-dividing states of the symbiont cells (Table S4; Fig. S1). Table S5 shows two-way ANOVA results on cell length comparisons between dividing symbiont cells and non-dividing symbiont cells from the symbiotic organs of all the insect species, which showed that both cell state and host species have statistically significant effects on cell length (both *P* < 0.001), indicating that, as expected, symbiont sizes tend to differ among different insect species. Table S6 summarizes *P* values of the post-hoc Tukey HSD test comparing the cell lengths of dividing symbiont cells and non-dividing symbiont cells from the symbiotic organs of all the insect species, which once again showed that symbiont sizes tend to differ significantly among different insect species.

### Cell division dynamics from exponential to stationary phases of *E. coli* proliferation

In an attempt to gain insight into the relationship between bacterial cell division and proliferation, we monitored the cell division dynamics of the cultivable free-living model bacterium *Escherichia coli* (Fig. 6; Table S7). In the exponential proliferation phases wherein bacterial densities are low and available nutrients are abundant, dividing cell ratios were as high as around 50% (Fig. 6A, B; Table S7). By contrast, in the stationary phases wherein bacterial densities attain a plateau and available nutrients become scarce, dividing cell ratios were down to 11% (Fig. 6A, C; Table S7). The bacterial density and the dividing cell ratios showed a clear negative correlation, or possibly an on-to-off biphasic relationship (Fig. 6D).

**FIG 6.**
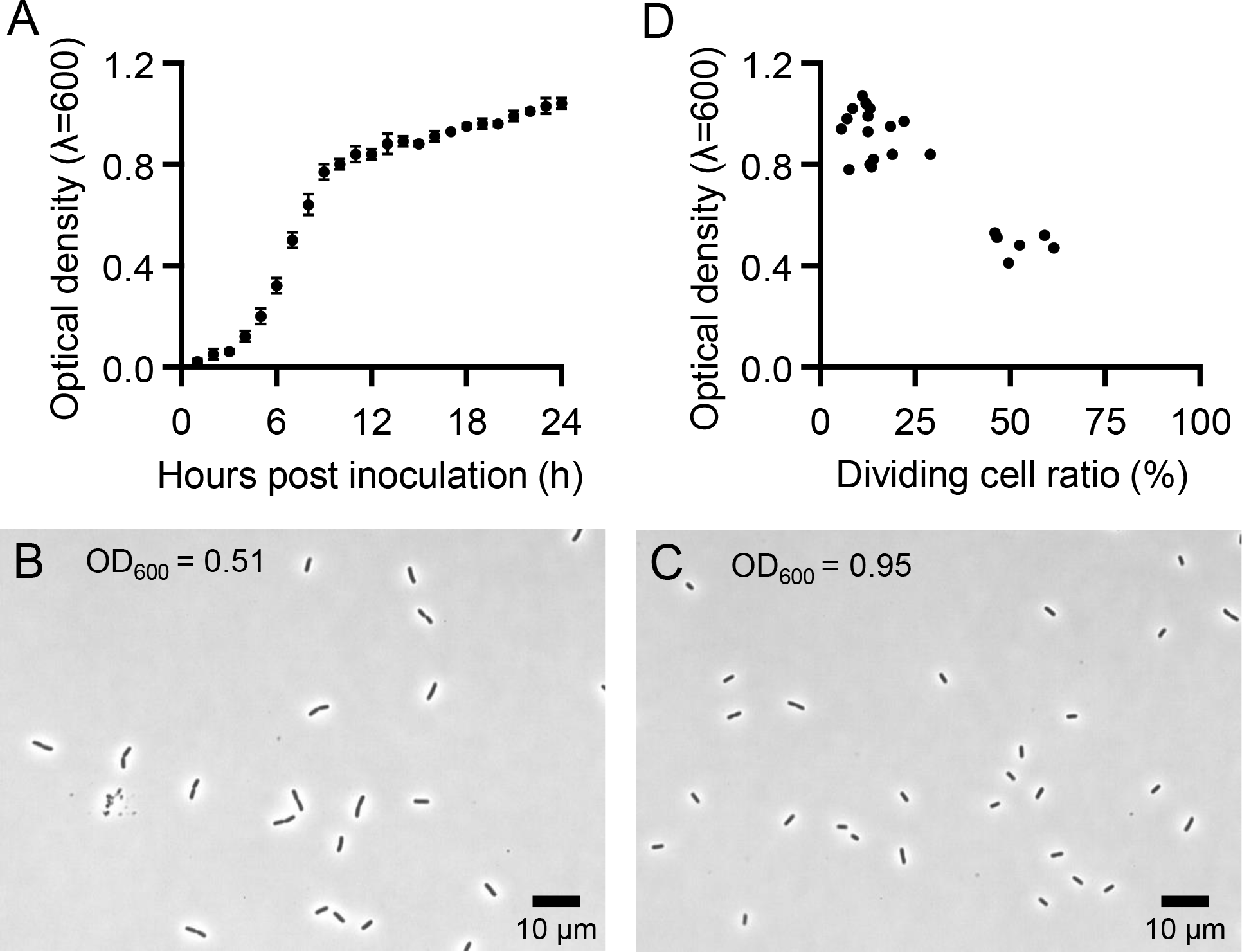
Cell division dynamics from exponential to stationary phases of *E. coli* proliferation. (**A**) Growth curve of *E. coli*, in which optical densities of replicate cultures (n = 4) are averaged and plotted with standard deviations. (**B**) Phase-contrast microscopic image of *E. coli* cells in an exponential phase of proliferation, in which the majority of cells are in division. (**C**) Phase-contrast microscopic image of *E. coli* cells in a stationary phase of proliferation, in which only a few cells are in division. (**D**) Relationship between *E. coli* proliferation and dividing cell ratio (n = 22). Also see Table S7.

### Comparative genomics of symbiont genes related to cell division

Finally, we surveyed the presence or absence of cell division-related bacterial genes involved in peptidoglycan biosynthesis, divisome formation, elongasome formation and peptidoglycan modification (37) among the genomes of *Blattabacterium* endosymbiont of the cockroach *B. germanica*, *Buchnera* endosymbiont of the aphid *A. pisum*, *Pantoea* gut symbiont of the stinkbug *P. stali*, and the laboratory model bacterium *E. coli* (Table S8). The cultivable free-living bacterium *E. coli*, whose genome is 4.6 Mb in size, retained almost all the cell division-related genes except for *mipZ*, a gene identified from *Caulobacter crescentus* whose product binds to FtsZ monomers to prevent their association with FtsZ filaments (38) and also caps the plus end of FtsZ filaments to promote their depolymerization (39). *mipZ* was also absent in the genomes of *Blattabacterium*, *Buchnera* and *Pantoea*, suggesting that it may be a gene specific to bacterial lineages allied to *C. crescentus.* The genome of the uncultivable stinkbug gut symbiont *Pantoea* sp. A, which is, though large in size (3.9 Mb), accumulating many pseudogenes and presumably at an evolutionarily early stage of obligatory symbiosis (40), retained the majority of the cell division-related genes. On the other hand, the reduced genomes of the cockroach endosymbiont *Blattabacterium* (0.64 Mb) and the aphid endosymbiont *Buchnera* (0.66 Mb) were devoid of some cell division-related genes in somewhat different patterns. Both symbiont genomes retained most of the peptidoglycan biosynthesis genes, reflecting the presence of cell wall in these endosymbiotic bacteria (16,17). Both symbiont genomes commonly lacked such divisome genes as *zipA*, *ftsE*, *ftsX* and *ftsK* that are involved in anchoring and activation of FtsZ filament to cell membrane (41–43), whereas *ftsL* and *ftsB* were absent only in *Blattabacterium* and *ftsQ* and *ftsN* were lacking only in *Buchnera*. Notably, the *Blattabacterium* genome was devoid of the core Min system genes *minC*, *minD* and *minE* that play a central role in positioning of Z-ring in bacterial cell division (44). It is also notable that the *Buchnera* genome lacked the Mre system genes *mreB*, *mreC*, *mreD* and *rodZ* that play an important role in bacterial cell elongation (45).

## DISCUSSION

In this study, we found that *Blattabacterium*, the endosymbiont of cockroaches, exhibits asymmetric cell division at a very high frequency. Since the first microscopic observation of the cockroach endosymbiont dating back to 19th century (46), some old hand-drawn sketches certainly depicted *Blattabacterium* cells showing asymmetric division (47,48), but little attention has been paid to the phenomenon for over a century. Needless to say, it is generally difficult to investigate the dynamics of cell division and proliferation of uncultivable symbiotic bacteria, but we obtained some insight into the challenging aspect of endosymbiosis by applying morphometric and statistical techniques to freshly prepared symbiont cells.

In the German cockroach *B. germanica*, the high frequency of asymmetric cell division of *Blattabacterium*, ranging from 28% to 58%, was consistently observed irrespective of host tissues and developmental stages (Figs. 1-3; Table S1). In other cockroach species like *P. japonica* and *E. yasumatsui*, the high frequency of asymmetric cell division of *Blattabacterium*, 35% and 24% respectively, was observed similarly (Fig. 4; Table S1), indicating that the frequent asymmetric division is a general trait of *Blattabacterium* endosymbionts of cockroaches. On the other hand, the asymmetric cell division was certainly observed in *P. indicus* but the frequency of cell division was low, only at 8% (Fig. 4; Table S1). The sizes of *Blattabacterium* cells were larger in adult fat bodies than in adult ovaries, nymphal ovaries and embryos irrespective of dividing or non-dividing cells (Fig. S1; Tables S2 and S4). These observations suggest that the peculiar traits of *Blattabacterium* may be affected or modified in physiological, developmental, ecological and/or evolutionary contexts. By contrast, in the aphid endosymbiont *Buchnera* and the stinkbug gut symbiont *Pantoea*, the cell division was symmetric binary fission typical of many bacteria (Fig. 5; Table S1), indicating that being symbiotic is not the major reason for asymmetric cell division. On the other hand, the *Buchnera* endosymbiont, which exhibits a drastically reduced genome size and uncultivability, shows a moderate frequency of cell division at 13%, whereas the *Pantoea* gut symbiont, which exhibits a moderately reduced genome size, eroded genes and uncultivability, shows a low frequency of cell division at 5% (Table S1). What mechanisms underlie these differences is currently elusive, but hereafter we attempt to discuss these issues in comparison with the model bacterium *E. coli* from viewpoints of cell division dynamics and comparative genomics.

When cell division and proliferation of *E. coli* was monitored during cultivation in a liquid medium, the cell division frequencies in the exponential phase were over 50% whereas the cell division frequencies in the stationary phase were down to 11% (Fig. 6). Hence, the cell division frequencies of *Blattabacterium* in the cockroaches are close to the levels at the exponential phase of *E. coli* when the bacterial cells actively divide, while the cell division frequencies of *Buchnera* in the aphid and *Pantoea* in the stinkbug are rather close to the levels at the stationary phase of *E. coli* when the bacterial growth is plateaued under nutritional and special restrictions. In the light of these observations, a possible explanation for the frequent cell division in *Blattabacterium* is that *Blattabacterium* is constantly in the exponential phase of cell proliferation. At a glance, this hypothesis may look strange because, densely packed in the cytoplasm of the host bacteriocytes, *Blattabacterium* seems to be under severe growth restriction. However, considering that the majority of ancient obligatory endosymbiotic bacteria, including *Blattabacterium*, have lost most regulatory genes in the course of reductive genome evolution (7,18), it is conceivable, although speculative, that cell division of *Blattabacterium* might be not regulated substantially but constantly ongoing at a very slow rate due to resource and space limitations. The relatively higher level of *Buchnera* cell division (13%) in comparison to *Pantoea* cell division (5%) (Table S1) might also be accounted for by the different levels of genome reduction (0.66 Mb vs. 3.9 Mb; see Table S8) and consequent loss of the control over cell division.

Besides the frequent cell division, the asymmetric cell division is a unique attribute of *Blattabacterium* that is not observed in *Buchnera*, *Pantoea*, *E. coli* and most other bacteria (Figs. 1-4). Why does only *Blattabacterium* exhibit asymmetric cell division? Our comparative genomic analysis focusing on cell division-related bacterial genes identified an important clue – the *Blattabacterium* genome is devoid of the core Min system genes *minC*, *minD* and *minE* that are involved in the positioning of the Z-ring in bacterial cell division (44). In *E. coli* and other bacteria, disruption of the Min system genes was reported to cause disturbed cell division, which often results in production of elongated cells and/or small DNA-free mini cells (49–52). It was also found that the *Buchnera* genome lacked the Mre system genes *mreB*, *mreC*, *mreD* and *rodZ* that are involved in bacterial cell elongation (45). In *E. coli* and other bacteria, disruption of the Mre system genes was reported to result in spherical bacterial cell shape (53–55). These genomic patterns suggest the possibility that the history of gene losses during the reductive genome evolution may have governed the cell morphology of such endosymbiont lineages as *Blattabacterium* and *Buchnera*.

All these results, observations and considerations taken together, we suggest that, although speculative, the frequent and asymmetric cell division observed in the ancient *Blattabacterium* endosymbiont of cockroaches is not an adaptive trait but rather an evolutionary consequence/byproduct of the genome erosion that generally proceeds in long-lasting and intimate host-symbiont associations (7,18). From this viewpoint, the dazzlingly diverse shapes and sizes of endosymbiotic bacteria of insects (13,20–24) may be integrated into a coherent picture in the context of reductive genome evolution. Whether such a frequent asymmetric cell division is found in ancient endosymbiotic bacteria of other insects is of great interest and to be pursued in future studies.

## MATERIALS AND METHODS

### Insect materials

A stock population of the German cockroach *Blattella germanica*, which was derived from around 100 individuals purchased from Sumika Technoservice Corporation, Takarazuka, Japan, was established in our laboratory at the National Institute of Advanced Industrial Science and Technology (AIST), Tsukuba, Ibaraki, Japan. Developmental staging of larvae was done as described previously (17). Stock populations of the Japanese cockroach *Periplaneta japonica* and the Indian cockroach *Pycnoscelus indicus* were established from about 10 wild caught individuals from Tsukuba, Ibaraki, Japan and Nago, Okinawa, Japan, respectively. A stock population of the azure cockroach *Eucorydia yasumatsui* was established from 1 wild caught individual from Ishigaki, Okinawa, Japan, which was provided by Shizuma Yanagisawa (Ryuyo Insect Nature Observation Park, Iwata, Japan). The insects were reared in plastic containers at 27°C under a 12 h light and 12 h dark regime in a climate chamber (MLR-352H-PJ, Panasonic) with insect feed (Insect Diet I, Oriental Yeast Co., Ltd.) and water. The pea aphid *Acyrthosiphon pisum* and the brown-winged green stinkbug *Plautia stali* used in this study were both laboratory strains established from adult insects collected in Tsukuba, Ibaraki, Japan, which were reared as described previously (32,56).

### Sample preparation and observation of symbiotic bacteria

The insects were dissected in a phosphate buffered saline (PBS: 0.8% NaCl, 0.02% KCl, 0.115% Na_2_HPO_4_; 0.02% KH_2_PO_4_ [pH 7.4]), and the isolated symbiotic organs (visceral fat bodies of the cockroaches, posterior midgut of the stinkbug, and bacteriocytes of the aphid) were placed in plastic tubes. The tissues were lightly agitated in PBS using a pestle to break the tissue apart while ensuring that the bacterial cells were not damaged. To remove uric acid crystals accumulated in the fat body cells of cockroaches and any other debris, the cell suspension was lightly centrifuged at low speeds and then passed through a 40 µm Flowmi Cell Strainer. The cell suspension was placed on a 22 x 40 mm coverslip and covered with an 18 x 18 mm coverslip, and then observed by phase-contrast microscopy using an inverted microscope (IX71; Olympus, Tokyo, Japan). These procedures were conducted as promptly as possible to minimize the damage to the fragile symbiont cells. Cell images were recorded by a complementary metal-oxide-semiconductor camera (DMK33UP5000; The Imaging Source, Bremen, Germany).

### Bacterial morphometry

Cell images were analyzed by ImageJ 1.54f. For cell length analysis, the major axis line of individual cells was traced and measured. For asymmetric level analysis, the cell length of the longer daughter cell was divided by that of the shorter daughter cell in each dividing cell. Plots were illustrated by IGOR Pro 8.04J (WaveMetrics, Portland, OR, United States).

### Statistical analysis

A two-way ANOVA was performed to analyze the effect of cell state (dividing/non-dividing) and the tissue type of *B. germanica* on bacterial cell length. A separate two-way ANOVA was performed to analyze the effect of cell state (dividing/non-dividing) and host species (including *P. stali* and *A. pisum*) on bacterial cell length. To determine any statistical difference between subgroups, a Tukey’s HSD test as well as independent Student’s t-tests were conducted for certain combinations. Both tests were done using R Studio version 2023.12.1.402 running version 4.0.3 (RStudio, PBC, Boston, MA).

### Electron microscopy

For backscattered electron scanning electron microscopy, fat bodies and ovaries were dissected from *B. germanica* adults in cold PBS. The samples were fixed overnight at 4°C in the mixture of 4% paraformaldehyde (Cat. No. 162-16065, Wako Pure Chemical Industries) and 2% glutaraldehyde (Sigma-Aldrich, G5882) in PBS, incubated with 2% osmium tetroxide in distilled water at 4°C for 2 h, dehydrated in a graded ethanol series (50%, 60%, 70%, 80%, 90%, 95%, 99%, and 100%), and treated with 100% propylene oxide followed by Epon 812. The treated samples were extracted by forceps and embedded in Epon 812. The hardened Epon blocks were sectioned using an ultramicrotome (ARTOS 3D, Leica) equipped with a diamond knife (SYM jumbo, 45 degrees, SYNTEK) to obtain 240 nm serial sections. The serial sections were collected on cleaned silicon wafer strips held by a micromanipulator (MN-153, NARISHIGE). The sections were stained at room temperature using 2% (w/v) aqueous uranyl acetate for 20 min and Reynolds’ lead citrate for 3 min. Images were obtained by a scanning electron microscope (JSM-7900F, JEOL).

### Culturing, preparation and observation of *E. coli*

*E. coli* strain BW25113 cells were statically cultured at 25°C in LB medium (5 g/L Yeast extract, 10 g/L Tryptone, 5 g/L NaCl). The 150 μL of *E. coli* pre-culture (OD_600_ = 0.4) was inoculated into 15 mL LB medium and statically cultured at 25°C. The OD_600_ of culture was measured by a spectrophotometer (GeneQuant1300; GE HealthCare, IL, United States) with 1-h intervals. Cultured cells at middle exponential phase and stationary phases were observed by phase-contrast microscopy and analyzed by ImageJ 1.54f.

### Comparative symbiont genomics

The genes related to bacterial cell division, which were selected according to Harpring and Cox (37), were obtained from the genome of *E. coli* K-12 (GenBank accession no. NC_000913.3). Then, orthologues of these genes were searched using the BLASTP program on CLC genomics workbench Ver. 24 (QIAGEN, Germany) against the genomes of *Blattabacterium* sp. strain Bge (GCF_000022605.2) (35), *Buchnera aphidicola* strain APS (GCF_000009605.1) (57) and stinkbug symbiont *Pantoea* sp. strain Ps-TKBA36 (GCF_001485275.1) (40). The resultant ortholog table was modified and verified manually with reference to KEGG orthology database (58).

## SUPPLEMENTARY MATERIALS

**FIG S1** Comparisons of the lengths of non-dividing symbiont cells from adult fat bodies, adult ovaries, nymphal ovaries and embryos, and also lengths of dividing symbiont cells from adult fat bodies, adult ovaries, nymphal ovaries and embryos. Different alphabetical letters (a, b) indicate statistically significant differences within the categories, namely non-dividing symbiont cells and dividing symbiont cells, respectively (Tukey HSD test: *P* < 0.05).

**TABLE S1** Dividing cell ratios of the symbiotic bacteria of cockroaches and other insects.

**TABLE S2** Sizes and ratios of dividing and non-dividing cells of the symbiotic bacteria of cockroaches and other insects.

**TABLE S3** Two-way ANOVA on cell length comparisons between dividing symbiont cells and non-dividing symbiont cells from different tissues of *B. germanica*.

**TABLE S4** Summary of *P* values of Tukey HSD test on cell length comparisons between dividing symbiont cells and non-dividing symbiont cells from different tissues of *B. germanica*. Bold shows statistically significant differences.

**TABLE S5** Two-way ANOVA on cell length comparisons between dividing symbiont cells and non-dividing symbiont cells from the symbiotic organs of all the insect species.

**TABLE S6** Summary of *P*-values of Tukey HSD test on cell length comparisons between dividing symbiont cells and non-dividing symbiont cells from the symbiotic organs of all the insect species. Bold shows statistically significant differences.

**TABLE S7** Dividing cell ratios of cultured *E. coli* cells.

**TABLE S8** Genes involved in peptideglycan biosynthesis, divisome, elongasome, and peptidoglycan modification encoded on the genomes of *E. coli*, *Blattabacterium*, *Buchnera* and *Pantoea*.

## ACKNOWLEDGMENTS

We thank Haruyasu Kohda and Keiko Okamoto-Furuta for their support in electron microscopic analysis. We also thank Shizuma Yanagisawa for providing insect samples.

This study was supported by the Japan Science and Technology Agency (JST) ERATO Grant to T.F., R.K. and T.H. (JPMJER1902), the Japan Society for the Promotion of Science (JSPS) Grant to T.H. (JP24H02294 and JP24K08935), and the Hakubi Project of Kyoto University to T.H. The Japan Society for the Promotion of Science (JSPS) Research Fellowships for Young Scientists supported T.N. (JP22KJ1191 and JP21J20814) and M.M. (JP22KJ3181 and JP22J00711).

## Author contributions

TN and MM equally contributed to this work. TN reared and collected all insects, processed the samples for microscopic observations, discovered the frequent asymmetric division of endosymbiont cells, and conducted post-hoc statistical analysis. MM conducted bacterial image acquisition and processing, and morphometric and statistical analyses. TH and TK performed electron microscopy. RK conducted comparative genomic analysis of cell division-related genes. TF conceived the research and wrote the paper. All authors contributed to and approved the final version of the manuscript.

## REFERENCES

1. Donachie WD. 1993. The cell cycle of *Escherichia coli*. Annu Rev Microbiol 47:199–231.

2. Adams DW, Errington J. 2009. Bacterial cell division: assembly, maintenance and disassembly of the Z ring. Nat Rev Microbiol 7:643–653.

3. Osella M, Tans SJ, Cosentino Lagomarsino M. 2017. Step by step, cell by cell: quantification of the bacterial cell cycle. Trends Microbiol 25:250–256.

4. Angert ER. 2005. Alternatives to binary fission in bacteria. Nat Rev Microbiol 3:214–224.

5. Young KD. 2006. The selective value of bacterial shape. Microbiol Mol Biol Rev 70:660–703.

6. Kysela DT, Randich AM. Caccamo PD, Brun YV. 2016. Diversity takes shape: understanding the mechanistic and adaptive basis of bacterial morphology. PLoS Biol 14:e1002565.

7. Moran NA, McCutcheon JP, Nakabachi A. 2008. Genomics and evolution of heritable bacterial symbionts. Annu Rev Genet 42:165–190.

8. Douglas AE. 2009. The microbial dimension in insect nutritional ecology. Funct Ecol 23:38–47.

9. Brune A. 2014. Symbiotic digestion of lignocellulose in termite guts. Nat Rev Microbiol 12:168–180.

10. Li H, Young SE, Poulsen M, Currie CR. 2021. Symbiont-mediated digestion of plant biomass in fungus-farming insects. Annu Rev Entomol 66:297–316.

11. Oliver KM., Smith AH, Russell JA. 2014. Defensive symbiosis in the real world– advancing ecological studies of heritable, protective bacteria in aphids and beyond. Funct Ecol 28:341–355.

12. Flórez LV, Biedermann PH., Engl T, Kaltenpoth M. 2015. Defensive symbioses of animals with prokaryotic and eukaryotic microorganisms. Nat Prod Rep 32:904–936.

13. Buchner P. 1965. Endosymbiosis of Animals with Plant Microorganisms. New York, NY: Interscience.

14. Salem H, Florez L, Gerardo N, Kaltenpoth M. 2015. An out-of-body experience: the extracellular dimension for the transmission of mutualistic bacteria in insects. Proc R Soc B 282:20142957.

15. Oguchi K, Harumoto T, Katsuno T, Matsuura Y, Chiyoda S, Fukatsu T. 2024. Intracellularity, extracellularity, and squeezing in the symbiotic organ underpin nurturing and functioning of bacterial symbiont in leaf beetles. iScience 27:109731.

16. Koga R, Meng XY, Tsuchida T, Fukatsu T. 2012. Cellular mechanism for selective vertical transmission of an obligate insect symbiont at the bacteriocyte–embryo interface. Proc Natl Acad Sci USA 109:E1230–E1237.

17. Noda T, Okude G, Meng XY, Koga R, Moriyama M, Fukatsu T. 2020. Bacteriocytes and *Blattabacterium* endosymbionts of the german cockroach *Blattella germanica*, the forest cockroach *Blattella nipponica*, and other cockroach species. Zool Sci 37:399–410.

18. McCutcheon JP, Moran NA. 2012. Extreme genome reduction in symbiotic bacteria. Nat Rev Microbiol 10:13–26.

19. Masson F, Lemaitre B. 2020. Growing ungrowable bacteria: Overview and perspectives on insect symbiont culturability. Microbiol Mol Biol Rev 84: e00089–20.

20. Moran NA, Tran P, Gerardo NM. 2005. Symbiosis and insect diversification: an ancient symbiont of sap-feeding insects from the bacterial phylum *Bacteroidetes*. Appl Environ Microbiol 71:8802–8810.

21. Nakabachi A, Yamashita A, Toh H, Ishikawa H, Dunbar HE, Moran NA, Hattori M. 2006. The 160-kilobase genome of the bacterial endosymbiont *Carsonella*. Science 314:267.

22. Login FH, Balmand S, Vallier A, Vincent-Monégat C, Vigneron A, Weiss-Gayet M, Rochat D, Heddi. 2011. Antimicrobial peptides keep insect endosymbionts under control. Science 334:362–365.

23. Hirota B, Okude G, Anbutsu H, Futahashi R, Moriyama M, Meng XY, Nikoh N, Koga R, Fukatsu T. 2017. A novel, extremely elongated, and endocellular bacterial symbiont supports cuticle formation of a grain pest beetle. mBio 8:10–1128.

24. Okude G, Koga R, Hayashi T, Nishide Y, Meng XY, Nikoh N, Miyanoshita A, Fukatsu T. 2017. Novel bacteriocyte-associated pleomorphic symbiont of the grain pest beetle *Rhyzopertha dominica* (Coleoptera: Bostrichidae). Zool Lett 3:13.

25. Komaki K, Ishikawa H. 1999. Intracellular bacterial symbionts of aphids possess many genomic copies per bacterium. J Mol Evol 48:717–722.

26. Woyke T, Tighe D, Mavromatis K, Clum A, Copeland A, Schackwitz W, Lapidus A, Wu D, McCutcheon JP, McDonald BR, Moran NA, Bristow J, Cheng JF. 2010. One bacterial cell, one complete genome. PLoS One 5:e10314.

27. Nakabachi A, Moran NA. 2022. Extreme polyploidy of *Carsonella*, an organelle-like bacterium with a drastically reduced genome. Microbiol Spectr 10:e00350–22.

28. Koga R, Tsuchida T, Fukatsu T. 2003. Changing partners in an obligate symbiosis: a facultative endosymbiont can compensate for loss of the essential endosymbiont *Buchnera* in an aphid. Proc R Soc B 270:2543-2550.

29. Wolschin F, Hölldobler B, Gross R, Zientz E. 2004. Replication of the endosymbiotic bacterium *Blochmannia floridanus* is correlated with the developmental and reproductive stages of its ant host. Appl Environ Microbiol 70:4096–4102.

30. Rio RV, Wu YN, Filardo G, Aksoy S. 2005. Dynamics of multiple symbiont density regulation during host development: tsetse fly and its microbial flora. Proc R Soc B 273:805–814.

31. Salem H, Bauer E, Kirsch R, Berasategui A, Cripps M, Weiss B, Koga R, Fukumori K, Vogel H, Fukatsu T, Kaltenpoth M. 2017. Drastic genome reduction in an herbivore’s pectinolytic symbiont. Cell 171:1520–1531.

32. Oishi S, Moriyama M, Koga R, Fukatsu T. 2019. Morphogenesis and development of midgut symbiotic organ of the stinkbug *Plautia stali* (Hemiptera: Pentatomidae). Zool Lett 5:16.

33. Lo N, Bandi C, Watanabe H, Nalepa C, Beninati T. 2003. Evidence for cocladogenesis between diverse dictyopteran lineages and their intracellular endosymbionts. Mol Biol Evol 20:907–913.

34. Sabree ZL, Kambhampati S, Moran NA. 2009. Nitrogen recycling and nutritional provisioning by *Blattabacterium*, the cockroach endosymbiont. Proc Natl Acad Sci USA 106:19521–19526.

35. López-Sánchez MJ, Neef A, Peretó J, Patiño-Navarrete R, Pignatelli M, Latorre A, Moya A. 2009. Evolutionary convergence and nitrogen metabolism in *Blattabacterium* strain Bge, primary endosymbiont of the cockroach *Blattella germanica*. PLoS Genet 5:e1000721.

36. Latorre A, Domínguez-Santos R, García-Ferris C, Gil R. 2022. Of cockroaches and symbionts: Recent advances in the characterization of the relationship between *Blattella germanica* and its dual symbiotic system. Life 12:290.

37. Harpring M, Cox JV. 2023. Plasticity in the cell division processes of obligate intracellular bacteria. Front Cell Infect 13:1205488.

38. Thanbichler M, Shapiro L. 2006. MipZ, a spatial regulator coordinating chromosome segregation with cell division in *Caulobacter*. Cell 126:147–162.

39. Corrales-Guerrero L, Steinchen W, Ramm B, Mücksch J, Rosum J, Refes Y, Heimerl T, Banger G, Schwille P, Thanbichler M. 2022. MipZ caps the plus-end of FtsZ polymers to promote their rapid disassembly. Proc Natl Acad Sci USA 119:e2208227119.

40. Hosokawa T, Ishii Y, Nikoh N, Fujie M, Satoh N, Fukatsu T. 2016. Obligate bacterial mutualists evolving from environmental bacteria in natural insect populations. Nat Microbiol 1:15011.

41. Barrows JM, Goley ED. 2021. FtsZ dynamics in bacterial division: What, how, and why? Curr Opin Cell Biol 68:163-172.

42. Du S, Lutkenhaus J. 2019. At the heart of bacterial cytokinesis: the Z ring. Trends Microbiol 27:781–791.

43. Yu XC, Tran AH, Sun Q, Margolin W. 1998. Localization of cell division protein FtsK to the *Escherichia coli* septum and identification of a potential N-terminal targeting domain. J Bacteriol 180:1296–1304.

44. Ramm B, Schumacher D, Harms A, Heermann T, Klos P, Müller F, Schwille P, Søgaard-Andersen L. 2023. Biomolecular condensate drives polymerization and bundling of the bacterial tubulin FtsZ to regulate cell division. Nat Commun 14:3825.

45. Busiek KK, Margolin W. 2015. Bacterial actin and tubulin homologs in cell growth and division. Curr Biol 25:R243–R254.

46. Blochmann F. 1887. Bakterienähnliche Körperchen in den Geweben und Eiern. Biol Zentralbl 7:606–608.

47. Gier HT. 1936. The morphology and behavior of the intracellular bacteroids of roaches. Biol Bull 71:433–452.

48. Koch A. 1949. Die Bakteriensymbiose der Küchenschaben. Mikrokosmos 38:121–125.

49. De Boer PA, Crossley RE, Rothfield LI. 1989. A division inhibitor and a topological specificity factor coded for by the minicell locus determine proper placement of the division septum in *E. coli*. Cell 56:641–649.

50. De Boer PA, Crossley RE, Rothfield LI. 1990. Central role for the *Escherichia coli* minC gene product in two different cell division-inhibition systems. Proc Natl Acad Sci USA 87:1129–1133.

51. De Boer PA, Crossley RE, Hand AR, Rothfield LI. 1991. The MinD protein is a membrane ATPase required for the correct placement of the *Escherichia coli* division site. EMBO J 10:4371–4380.

52. Bramkamp M, Emmins R, Weston L, Donovan C, Daniel RA, Errington J. 2008. A novel component of the division-site selection system of *Bacillus subtilis* and a new mode of action for the division inhibitor MinCD. Mol Microbiol 70:1556–1569.

53. Wachi M, Doi M, Okada Y, Matsuhashi M. New *mre* genes *mreC* and *mreD*, responsible for formation of the rod shape of *Escherichia coli* cells. J Bacteriol 171:6511–6516.

54. Jones LJ, Carballido-López R, Errington J. 2001. Control of cell shape in bacteria: helical, actin-like filaments in *Bacillus subtilis*. Cell 104:913–922.

55. Figge RM, Divakaruni AV, Gober JW. 2004. MreB, the cell shape-determining bacterial actin homologue, co-ordinates cell wall morphogenesis in *Caulobacter crescentus*. Mol Microbiol 51:1321–1332.

56. Fukatsu T, Nikoh N, Kawai R, Koga R. 2000. The secondary endosymbiotic bacterium of the pea aphid *Acyrthosiphon pisum* (Insecta: Homoptera). Appl Environ Microbiol 66:2748–2758.

57. Shigenobu S, Watanabe H, Hattori M, Sakaki Y, Ishikawa H. 2000. Genome sequence of the endocellular bacterial symbiont of aphids *Buchnera* sp. APS. Nature 407:81–86.

58. Kanehisa M, Furumichi M, Sato Y, Kawashima M, Ishiguro-Watanabe M. 2023. KEGG for taxonomy-based analysis of pathways and genomes. Nucleic Acids Res 51:D587– D592.

